# A high fat anti-inflammatory diet improves widespread allodynia despite worsening metabolic outcomes in adult mice exposed to neonatal maternal separation

**DOI:** 10.1101/2020.09.29.317297

**Authors:** Olivia C. Eller, Rebecca M. Foright, Aaron D. Brake, Michelle K. Winter, Leonidas E. Bantis, E. Matthew Morris, John P. Thyfault, Julie A. Christianson

## Abstract

Inflammation plays a key role in the progression and maintenance of chronic pain, which impacts the lives of millions of Americans. Despite growing evidence that chronic pain can be improved by treating underlying inflammation, successful treatments are lacking and pharmaceutical interventions are limited due to drug side effects. Here we are testing whether an anti-inflammatory diet (AID) containing a combination of key anti-inflammatory compounds, at clinically relevant doses, improves pain-like behaviors in a preclinical model of chronic widespread hypersensitivity induced by neonatal maternal separation (NMS). Our results demonstrate a benefit of the AID on pain-like behaviors, despite the diet being high in fat, which led to increased caloric intake, adiposity, and weight gain. The AID specifically increased measures of metabolic syndrome and inflammation in female mice, compared to an isocaloric, macronutrient-matched diet lacking the anti-inflammatory compounds. Male mice, especially those exposed to NMS, were equally susceptible to both diets worsening metabolic measures. This work highlights important sexual dimorphic outcomes related to early life stress exposure and dietary interventions, as well as a potential disconnect between improvements in pain-like behaviors and metabolic measures.

## Introduction

Chronic pain conditions affect 40-116 million Americans (1, 2), costing an estimated $635 billion annually in health care costs and lost work productivity (3). Sustained, low-grade inflammation is thought to be a contributing factor in many chronic pain disorders (4). Unresolved inflammation following an acute injury can contribute to the development of chronic pain through the continued release of prostaglandins, bradykinin, and proinflammatory cytokines and chemokines that activate nociceptors and drive peripheral sensitization (5–8). Increased visceral fat accumulation, a key hallmark of obesity and metabolic disorder, also leads to chronic low-grade inflammation (9) and has been postulated to underlie neuroimmune mechanisms that contribute to chronic pain (10). This is clinically important as obesity and chronic pain are known comorbid disease conditions, despite a paucity of work examining if these conditions are mechanistically linked. Pharmacological-based treatment of the inflammatory component of chronic pain in either lean or obese states has been limited due to harmful side effects (11, 12), which has led to research on the effectiveness of diet-based interventions to lower inflammation (13).

Generally, an anti-inflammatory diet (AID) is euglycemic and high in fruits, vegetables, lean protein, whole grains high in fiber, healthy fats such as those high in omega-3 fatty acids, and/or supplementation with anti-inflammatory compounds (14). Supplemental compounds include epigallocatechin gallate (EGCG), sulforaphane, resveratrol, curcumin, and ginseng. EGCG is the most abundant polyphenol found in green tea (15) and has been shown to increase mechanical and thermal pain thresholds in a model of chronic constriction injury (CCI) (16). Sulforaphane is found in cruciferous vegetables (17, 18) and, like EGCG, prevented the development of CCI-induced mechanical and thermal hypersensitivity (19). Resveratrol is a natural polyphenol and phytoalexin with anti-inflammatory and antioxidant effects (20) and was shown to attenuate thermal hyperalgesia in a mouse model of diabetic neuropathy (21). Curcumin is a bioactive polyphenol found in turmeric (22). Clinically, curcumin has been effective in the treatment of knee osteoarthritis (23) and preclinically, prevented the development of thermal and mechanical paw hypersensitivity in CCI (24). Ginseng is a root that contains pharmacological compounds called ginsenosides (25) and prevented pain-like behaviors following capsaicin injection into the hindpaw (26). There is growing evidence that these individual anti-inflammatory compounds can improve aspects of chronic pain, however, these studies are limited (16, 19, 21, 23, 24, 26, 27) and additional studies using a combination of these compounds at physiologically relevant doses are warranted (28).

In the present study, we determined the effectiveness of an AID on attenuating outcomes associated with early life stress in our mouse model of neonatal maternal separation (NMS). The NMS model exhibits evidence of persistent low-level inflammation displayed as urogenital hypersensitivity associated with local neuroimmune activation (29–32), altered glucocorticoid production and receptor expression within the limbic system and hypothalamic-pituitary-adrenal (HPA) axis (29–32), and increased susceptibility to obesity on both a chow and high fat/high sucrose diet (33). The AID used in the present study, developed from Totsch et. al., (28), contained EGCG, sulforaphane, resveratrol, curcumin, ginseng and fats with a high omega-3 to omega-6 fatty acid ratio. While we found that the AID effectively reduced NMS-induced urogenital and widespread hypersensitivity, the diet negatively impacted weight gain and adiposity, as well as metabolic health.

## Methods

### Animals

All experiments were performed on male and female C57Bl/6 mice (Charles River, Wilmington, MA) born and housed in the Research Support Facility at the University of Kansas Medical Center. Mice were housed at 22° C on a 12-hour light cycle (600 to 1800 hours) and received water and food *ad libitum*. All research was approved by the University of Kansas Medical Center Institutional Animal Care and Use Committee in compliance with the National Institute of Health Guide for the Care and Use of Laboratory Animals.

### Neonatal maternal separation (NMS)

Pregnant C57Bl/6 dams were delivered to the animal facility between 14-16 days gestation. Litters were divided equally into NMS and naïve groups. From postnatal day 1 (P1) until P21, NMS pups were removed *en masse* and placed in a clean glass beaker with bedding from their home cage for 180 minutes (11am-2pm). The beaker was placed in an incubator maintained at 33°C and 50% humidity. Naïve mice remained undisturbed in their home cage except for normal animal husbandry. All mice were weaned on P22 and pair-housed with same sex litter mates and *ad libitum* access to water and a control diet composed of 20% kcal protein, 70% kcal carbohydrate (3.5% sucrose), and 10% kcal fat (Research Diets, Inc. New Brunswick, NJ, D17072402; Table 1).

### Anti-inflammatory diet (AID)

At 4 weeks of age, female and male naïve and NMS mice were further divided. Half remained on the control diet and half were given an AID, which was composed of 20% kcal protein, 45% kcal carbohydrate (0% sucrose), and 35% kcal fat with a Hi-Maize 260 starch source and added anti-inflammatory compounds: EGCG, sulforaphane, resveratrol, curcumin, and ginseng (Research Diets, Inc. New Brunswick, NJ, D17072401; Table 1).

### Non-anti-inflammatory diet (NAID)

In a different cohort of mice, at 4 weeks of age female and male naïve and NMS mice were further divided. Half remained on the control diet and half were given the NAID, which had the same caloric and macronutrient breakdown of the AID without the added anti-inflammatory components (Research Diets, Inc. New Brunswick, D17072403, NJ; Table 1).

### Perigenital mechanical sensitivity

Perigenital mechanical withdrawal threshold was assessed every 4 weeks starting after 4 weeks on the AID or NAID. For 2 days prior to the test day, mice were acclimated to a soundproof room for 30 minutes and then placed into individual clear plastic chambers (11 x 5 x 3.5 cm) on a wire mesh screen elevated 55 cm above a table for 30 minutes. Additionally, the perivaginal area of female mice was shaved on the first day of acclimation. On the test day, mice were acclimated to the soundproof room for 30 minutes and then placed on the table for 30 minutes. The up-down method was performed to test mechanical sensitivity using a standard set of Semmes-Weinstein monofilaments (1.65, 2.36, 2.83, 3.22, 3.61, 4.31, 4.74 g; Stoelting, Wood Dale, IL) (34, 35). Beginning with the 3.22 g monofilament, mice received a single application to either the scrotum or perivaginal area. A negative response was followed by the next larger filament and a positive response (considered a brisk jerk or licking the probed area) was followed by the next smaller filament. The experimenter continued to move up or down the series, depending on the previously elicited response, for an additional four applications after the first positive response was observed for a minimum of five or a maximum of nine total monofilament applications. The value in log units of the final monofilament applied in the trial series was used to calculate 50% g threshold for each mouse (34).

### Hindpaw mechanical sensitivity

Hindpaw mechanical sensitivity was assessed every 4 weeks starting after 5 weeks on the AID or NAID. On the test day, mice were acclimated to a soundproof room for 30 minutes and then placed into individual clear plastic chambers (11 x 5 x 3.5 cm) on a wire mesh screen elevated 55cm above a table for 30 minutes. An electronic von Frey device (IITC Life Science Inc. Woodland Hills, CA) was used to measure hindpaw withdrawal threshold. A semi-flexible tip filament was applied to the hindpaw and the force that elicited a withdrawal was recorded from the electronic device. The filament was applied 6 times to each mouse and the highest and the lowest value for each mouse were excluded. Therefore, an average of 4 measurements/mouse was quantified.

### Nest building test

At 1 hour before the start of the dark phase (5pm), mice were individually placed into clean cages containing no environmental enrichment outside of a 3g nestlet square. Seventeen hours later (10am), the mice were returned to their home cages and the nests were photographed and intact nestlet pieces were weighed. Two blinded experimenters scored the nests based on a 1 to 5 scale according to previous publications (36, 37) and the average of their scores is reported here.

### Body weight, intake and feed efficiency

Energy intake (per pair) and body weight were measured weekly. Feed efficiency was calculated as weight gained/calories consumed per pair of mice per week. An average feed efficiency was quantified from weekly feed efficiency.

### Body composition analysis

Every 4 weeks, mice were weighed and placed in an EchoMRI 2015 (EchoMRI LLC, Houston, TX) to quantify lean mass and fat mass. At time of euthanization, mice were overdosed with inhaled isoflurane, the epididymal/periovarian and retroperitoneal fat pads were excised and weighed. These two fat pad weights were summed to calculate total visceral fat.

### Fasting insulin

After 18-19 weeks on the AID or NAID, fasting insulin level was measured. Following a 6-hour fast, blood was collected via tail-clip, placed on ice for 1 hour, and centrifuged at 10,000 rpm for 10 minutes. Serum was collected and frozen until analysis using an insulin *ELISA* kit (80-INSMS-E01, ALPCO, Salem, NH) according to the manufacturer’s instructions.

### Glucose Tolerance Test

After 18-19 weeks on the AID or NAID, a glucose tolerance test was carried out. Following a 6-hour fast, mice were given an IP injection of glucose at 1 g/kg body weight. Blood was collected via tail clip immediately prior to the glucose injection and 15, 30, 60, and 120 minutes thereafter and blood glucose concentrations were measured by colorimetric assay (PGO enzyme preparation and dianisidine dihydrochloride, Sigma-Aldrich, St. Louis, MO).

### HOMA-IR

To calculate HOMA-IR we used the formula: fasting insulin (mU/L) * fasting glucose (mg/dL)/405.

### Corticosterone

After 18-19 weeks on the AID or NAID, mice were sacrificed during the early half of the light cycle (8:00am −11:00am) and trunk blood was collected. Serum was removed and frozen until analysis using a corticosterone ELISA kit (55-CORMS-E01, ALPCO, Salem, NH) according to the manufacturer’s instructions.

### mRNA extraction and RT-PCR

Epidydimal and periovarian adipose tissue was dissected, weighed, and immediately frozen in liquid nitrogen, and stored at −80°C. Frozen tissue was then crushed (Cellcrusher, Portland, OR) and total RNA was isolated using QIAzol Lysis Reagent and the RNeasy Lipid Tissue Mini Kit (Qiagen, Valencia, CA). The concentration and purity were determined using NanoDrop 2000 (Thermo Fisher Scientific, Wilmington, DE) and cDNA was synthesized from total RNA using the iScript cDNA Synthesis Kit (Bio-Rad, Hercules, CA). Quantitative RT-PCR was performed using SsoAdvanced SYBR Green Supermix (Bio-Rad) and a Bio-Rad iCycler IQ real time PCR system with indicated 20uM primers (Table 2; Integrated DNA Technologies, Coralville, IA). To reduce variability due to fluctuations in baseline fluorescence, the raw PCR data was imported to the LinRegPCR software and PCR efficiency values were derived for each individual sample. Threshold cycle values were subtracted from that of the housekeeping gene PPiB and the fold change over Naïve-Control was calculated using the Pfaffl method (38)

### Statistical analyses

Comparisons were made between the control groups from the AID and NAID cohorts based on sex and NMS status and no significant differences were observed for perigenital sensitivity, hindpaw sensitivity, nest building, body weight, body fat, food intake, feed efficiency, fasting insulin, serum glucose levels, serum corticosterone, or RT-PCR; therefore, the control groups were combined for statistical comparisons. Calculations were made in Excel (Microsoft, Redmond, WA) and normality of distribution was tested using Shapiro-Wilk’s test (*p*<0.05). Non-normally distributed data were log transformed prior to statistical analysis. Statistical analyses were performed using GraphPad Prism 8 (GraphPad, La Jolla, CA) or IBM SPSS Statistics 25 (IBM Corporation, Armonk, NY). Differences between groups were determined by two- or three-way ANOVA, with or without repeated measures (RM) and Fisher’s LSD posttest. Statistical significance was set at *p*<0.05. Statistical analyses of mechanical sensitivity were validated by using a Generalized Estimating Equation (GEE) framework. For hindpaw sensitivity, the following model was created for each diet/sex: 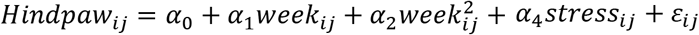 and for perigenital sensitivity the following model was created for each diet/sex: 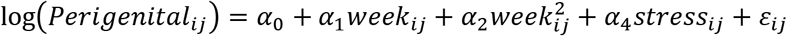. To fit these models, a Markov structure was employed for the working correlation matrix. For posttest analyses, each subgroup was compared in terms of area under the curve.

## Results

### AID reduces perigenital mechanical sensitivity

Perigenital mechanical sensitivity was measured every 4 weeks and overall sensitivity changes across the experiment were calculated as area under the curve (Figure 1). A significant effect of diet was observed on perigenital withdrawal threshold in the female mice (Figure 1A-B). In both naïve and NMS female mice, the AID resulted in a significantly higher perigenital withdrawal threshold AUC compared to control-fed and NAID-fed mice, indicating that AID reduced perigenital mechanical sensitivity (Figure 1B). In male mice, there was an overall effect of diet on perigenital withdrawal threshold (Figure 1C-D), with AID-fed naïve and NMS male mice both exhibiting significantly higher mechanical withdrawal thresholds compared to control-fed and NAID-fed mice. In addition, NMS-NAID male mice had significantly lower withdrawal thresholds compared to NMS-control (Figure 1D). Taken together, these data suggest that AID reduced perigenital mechanical sensitivity in all mice, regardless of sex or NMS exposure, while NAID selectively increased perigenital mechanical sensitivity only in NMS male mice.

**Figure 1.**
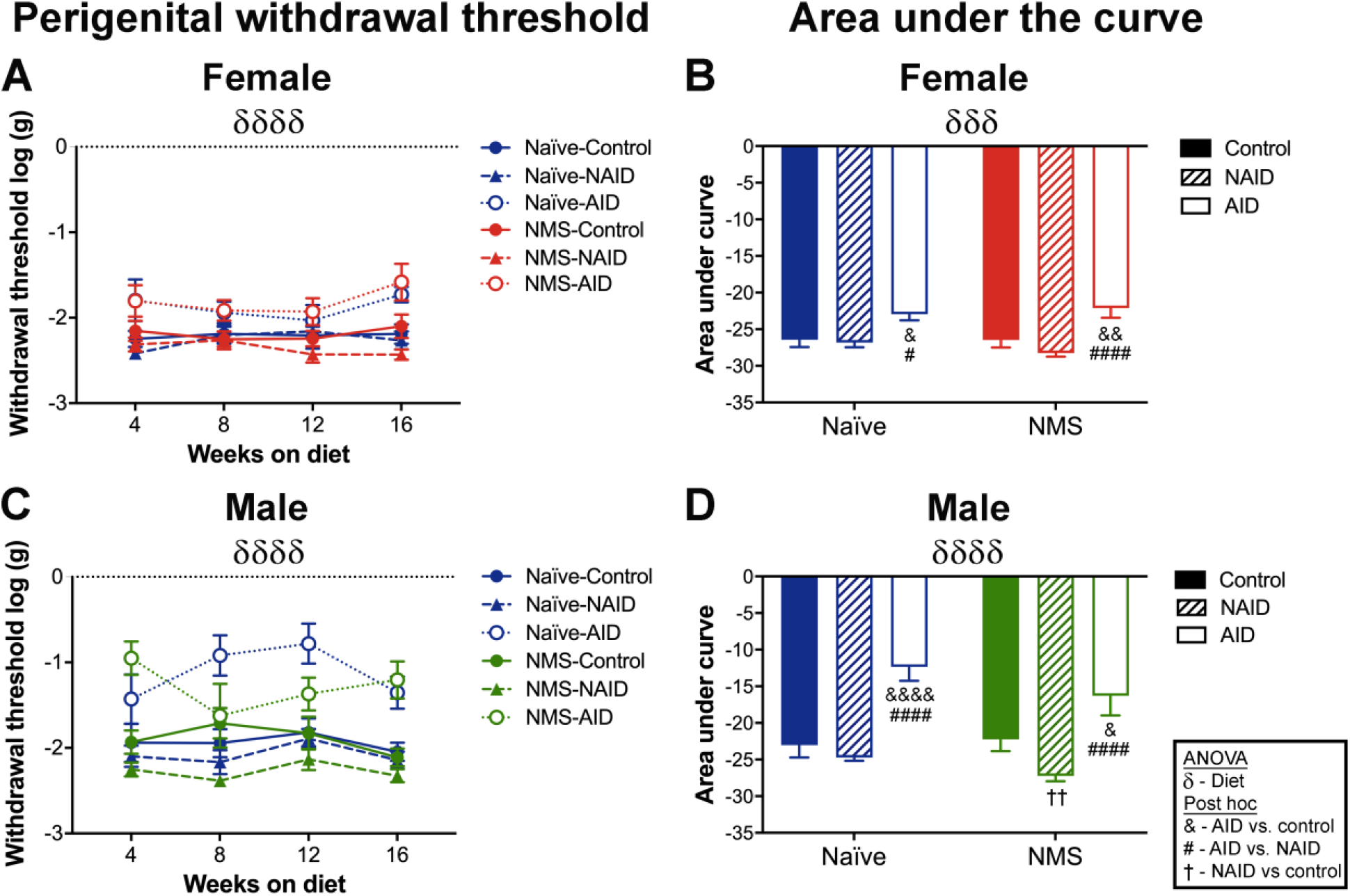
Impact of NMS and diet on perigenital mechanical withdrawal threshold. Perigenital withdrawal threshold was measured every 4 weeks in female (**A**) and male (**C**) mice and the area under the curve (AUC) was calculated (**B, D**). **A**) In female mice, there was a significant effect of diet over the 16 weeks (*p*<0.0001). **B**) AUC measurements also revealed a significant effect of diet (*p*=0.0005) with AID-fed mice having significantly higher thresholds than NAID- or control-fed mice, regardless of NMS status. **C**) In males, there was a significant effect of diet (*p*<0.0001) over the 16 weeks. **D**) AUC measurements also revealed a significant effect of diet (*p*<0.0001) with AID-fed mice having significantly lower withdrawal thresholds compared to NAID- and control-fed for both naïve and NMS mice. Additionally, NMS-NAID mice had significantly greater withdrawal thresholds than both NMS-Control and Naïve-NAID mice. δ denotes a significant effect of diet, three-way RM ANOVA (**A, C**) or two-way ANOVA (**B, D**). &, &&, &&&& *p*<0.05, 0.01, 0.0001 AID vs. control, #, #### *p*<0.05, 0.0001 AID vs. NAID, †† *p*<0.01 NAID vs. control, Fisher’s LSD. n=8-16 per group.

### AID attenuates hindpaw allodynia in female NMS mice

In female mice, there was a significant impact of NMS on hindpaw mechanical withdrawal threshold both across all experimental time points (Figure 2A) and for the AUC measurements (Figure 2B). Female NMS-control and -NAID mice had a significantly lower hindpaw withdrawal threshold AUC compared to naïve-control and -NAID mice, respectively (Figure 2B). NMS-AID mice had a modest increase in hindpaw withdrawal threshold that was not significantly different from either naïve-AID, NMS-control, or NMS-NAID mice (Figure 2B). In male mice, there was a trend toward an NMS effect on hindpaw withdrawal thresholds both across the experimental time points (*p*=0.081, Figure 2C) and for the AUC measurements (*p*=0.079, Figure 2D). No effect of diet was observed on hindpaw mechanical withdrawal threshold, regardless of sex or NMS status. Together, these data suggest that NMS had a negative impact on hindpaw sensitivity, which was modestly improved by AID only in the female mice.

**Figure 2:**
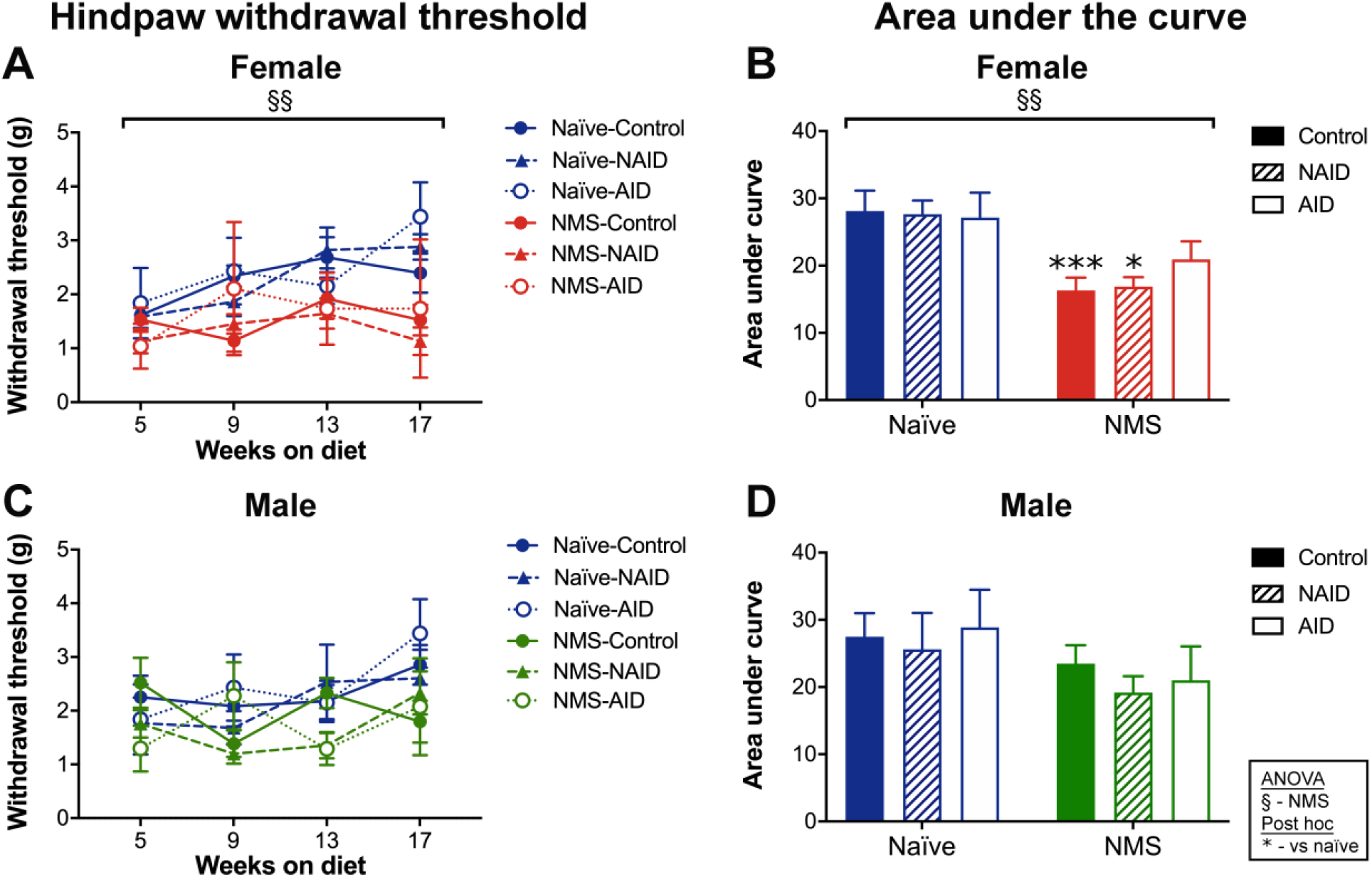
Impact of NMS and diet on hindpaw mechanical withdrawal threshold. Hindpaw mechanical withdrawal threshold was measured every 4 weeks (**A, C**) and the area under the curve (AUC, **B, D**) was calculated. In female mice, there was an overall significant effect of NMS across all experimental time points (**A**, *p*=0.001) and for the AUC measurement (**B**, *p*=0.001). **B**) Both NMS-Control and NMS-NAID female mice had significantly lower withdrawal thresholds compared to their naïve counterparts. **C, D**) In males, there was a trend toward an NMS effect, but no effect of diet. § denotes a significant impact of NMS, three-way RM ANOVA (**A**) or two-way ANOVA (**B**). *, *** *p*<0.05, 0.001 vs. same-diet naïve, Fisher’s LSD posttest. n=8-16 per group.

### AID worsened nest building scores in female NMS mice

Nest building was evaluated at the end of the experiment to determine the effect of NMS and diet on a measure of anhedonia. In the females, there was a significant impact of diet on nest scores (Figure 3A) and NMS-AID mice had significantly worse scores, compared to naïve-control mice. There were no observed NMS or diet impacts on nest building in male mice (Figure 3B). These data suggest that AID in female NMS mice results in anhedonic behavior.

**Figure 3.**
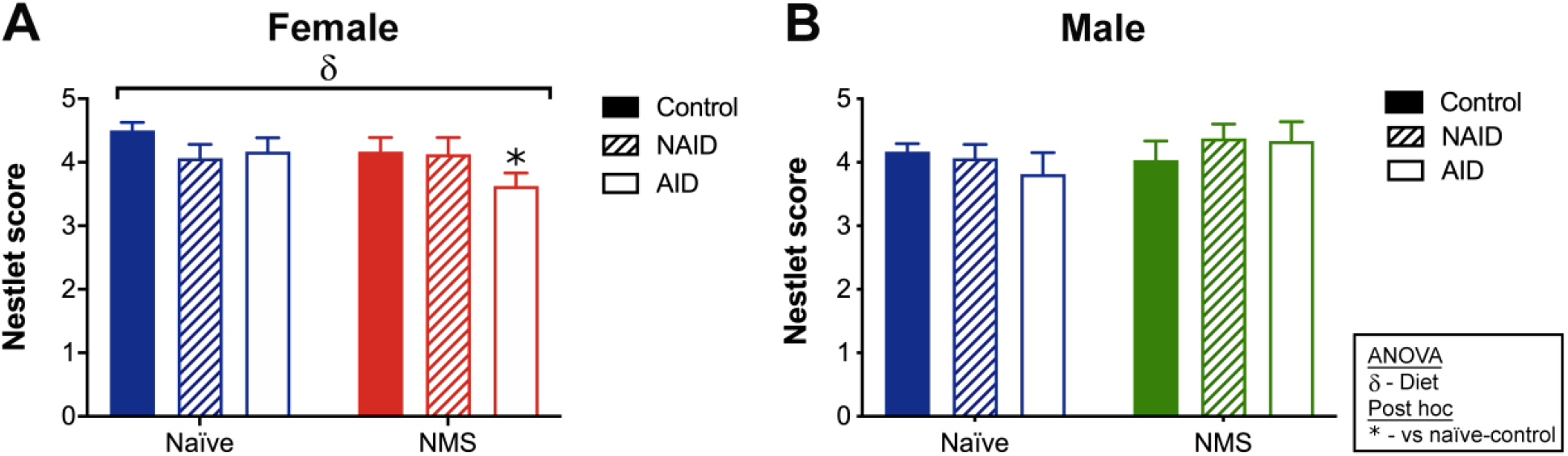
Impact of NMS and diet on nest building behavior. **A**) In female mice, there was a significant overall effect of diet (*p*=0.048) on nest quality. NMS-AID mice had a significantly lower nest score compared to naïve-control mice. **B**) Nest quality was not affected by diet or NMS in male mice. δ denotes significant effect of diet, two-way ANOVA. \**p*<0.05 vs. naïve-control mice, Fisher’s LSD posttest. n=8-16 per group.

### AID increased body weight and fat gain in female and male NMS mice

Body weight was measured weekly and body fat percentage was measured every 4 weeks throughout the course of the experiment (Figure 4). In females, there was a significant impact of NMS and diet on body weight across the entire experiment (Figure 4A) and at time of euthanization (Figure 5A). After 18 weeks on the diets, NAID- and AID-fed female mice were significantly heavier than control-fed, regardless of NMS status (Figure 5A). In addition, NMS-AID female mice were significantly heavier than naïve-AID mice. Only diet had a significant impact on body fat percentage across all time points (Figure 4B) and at euthanization (Figure 5B). Naïve-AID and NMS-AID mice had significantly higher body fat percentage compared to naïve-control and NMS-control mice, respectively (Figure 5B). NMS-AID mice also had a significantly higher body fat percentage compared to NMS-NAID mice (Figure 5B). Periovarian fat pads weight were significantly impacted by diet and were significantly heavier in NMS-AID female mice compared to either NMS-control or NMS-NAID mice (Figure 5C).

**Figure 4.**
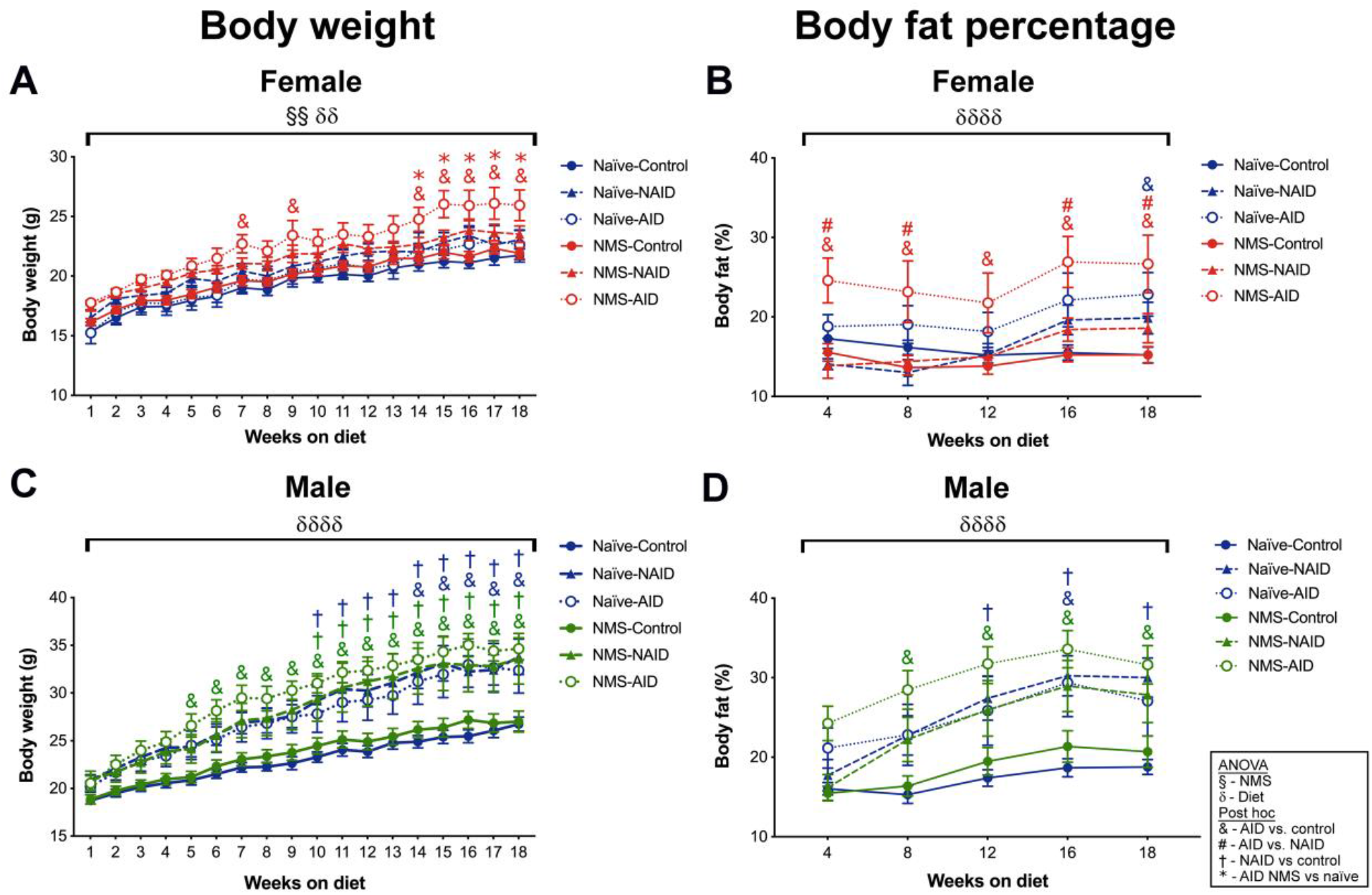
Impact of NMS and diet on body weight and composition. **A**) In female mice, there was a significant overall effect of NMS (*p*=0.007) and diet (*p*=0.006) on body weight across 18 weeks. NMS-AID mice weighed significantly more than NMS-control (weeks 7, 9, and 14-18) and NMS-NAID (weeks 14-18) mice. **B**) Body composition in females was significantly impacted by diet (*p*<0.0001). NMS-AID mice had significantly higher body fat percentage compared to NMS-control mice (every timepoint) and NMS-NAID mice (all but 12 weeks). At 18 weeks on the diet, naïve-AID mice had significantly higher body fat percentage than naïve-control mice. **C**) In male mice, there was a significant overall effect of diet (*p*<0.0001) on body weight. NMS-AID (weeks 5-18) and -NAID (weeks 10-18) mice weighed more than NMS-control mice. Similarly, naïve-AID (weeks 14-18) and -NAID (weeks 10-18) mice weighed more than naïve-control mice. **D**) Body composition analyses in male mice found a significant overall effect of diet (*p*<0.0001) on body fat percentage. NMS-AID mice had significantly higher body fat percentage compared to NMS-control mice (weeks 8-18). At week 16, naïve-AID mice had significantly higher body fat percentage compared to naïve-control mice and naïve-NAID had significantly higher body fat percentage compared to naïve-control (weeks 12-18). § and δ denote significant effects of NMS and diet, respectively, three-way RM ANOVA. \**p*<0.05 vs. same-diet naïve, & *p*<0.05 AID vs. control, † *p*<0.05 NAID vs. control, # *p*<0.05 AID vs. NAID, Fisher’s LSD posttest. n=8-16 per group.

**Figure 5.**
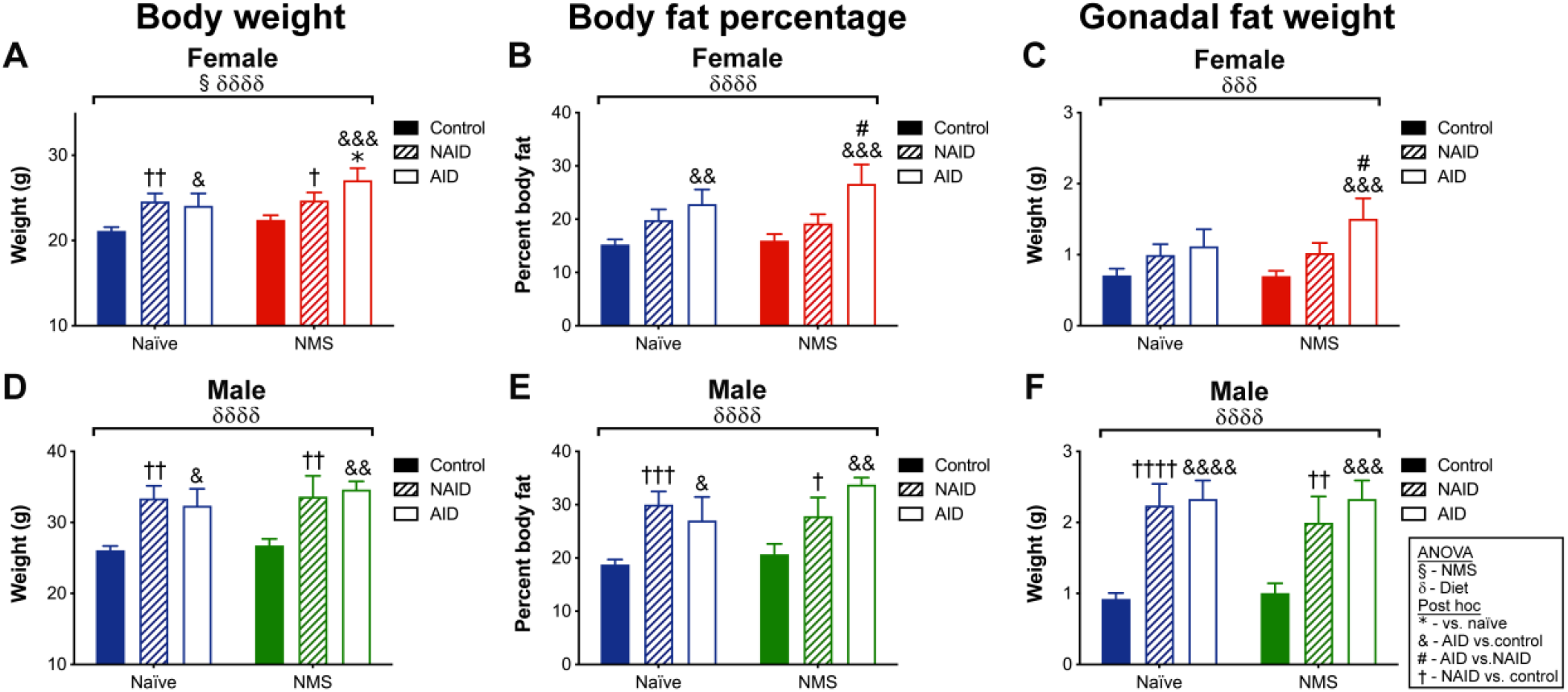
The impact of NMS and diet on end-of-study body weight, body fat percentage, and gonadal fat weight. **A)** Final body weight in female mice was significantly impacted by NMS (*p*=0.044) and diet (*p*<0.0001). NAID- and AID-fed mice were significantly heavier than control-fed mice, regardless of NMS status. NMS-AID female mice were also significantly heavier than naïve-AID mice. **B)** Final body fat percentage was significantly impacted by diet (*p*<0.0001), specifically in AID-fed mice. **C)** Periovarian fat weight was significantly impacted by diet (*p*<0.0001) with NMS-AID mice having significantly heavier fat pads than NMS-control and NMS-NAID mice. In male mice, final body weight **(D)**, body fat percentage **(E)**, and epididymal fat pad weight **(F)** were all significantly impacted by diet (*p*<0.0001) with NAID- and AID-fed mice having significantly higher values compared to control-fed mice, regardless of NMS status. § and δ denote significant effects of NMS and diet, respectively, two-way ANOVA. \**p*<0.05 vs. same-diet naïve, &, &&, &&&, &&&& *p*<0.05, 0.01, 0.001, 0.0001 AID vs. control, †, ††, †††, †††† *p*<0.05, 0.01, 0.001,0.0001 NAID vs. control, # *p*<0.05 AID vs. NAID, Fisher’s LSD posttest. n=8-16 per group.

In male mice, only diet significantly impacted body weight (Figure 4C) and body fat percentage (Figure 4D) across all time points. After 18 weeks on the diets, male NAID- and AID-fed mice had significantly greater body weight (Figure 5D) and body fat percentage (Figure 5E) compared to control-fed mice, regardless of NMS status. Epididymal fat pad weight was significantly impacted by diet and was significantly heavier in NAID- and AID-fed male mice compared to control-fed, regardless of NMS status (Figure 5F). Together these data suggest that the AID had a greater effect on driving adiposity in NMS female mice, whereas both NAID and AID drove weight and fat gains in naïve and NMS male mice to a similar extent.

### NAID and AID impacted food intake and feed efficiency

Food intake was measured weekly and is reported as calorie intake consumed per pair (Figure 6A, C). In females, calories consumed were significantly impacted by diet across the duration of the experiment (Figure 6A). Female naïve and NMS mice fed either AID or NAID consumed more calories/pair compared to control-fed mice at nearly every time point measured. Feed efficiency in female mice was also significantly impacted by diet, however, despite being equal in caloric density and similarly impacting caloric intake, AID-fed female mice had significantly increased feed efficiency compared to NAID-fed mice, regardless of NMS status (Figure 6B). In males, caloric intake was significantly impacted by diet (Figure 6C) with AID- and NAID-fed mice consuming significantly more calories than control-fed mice at nearly every time point, regardless of NMS status. Feed efficiency in male mice was also significantly impacted by diet, however, unlike in the female mice, there was no significant difference in feed efficiency between NAID- and AID-fed mice, regardless of NMS status (Figure 6D). These data suggest that AID specifically increases caloric intake and feed efficiency in female mice, while both NAID and AID similarly impact these measures in male mice.

**Figure 6.**
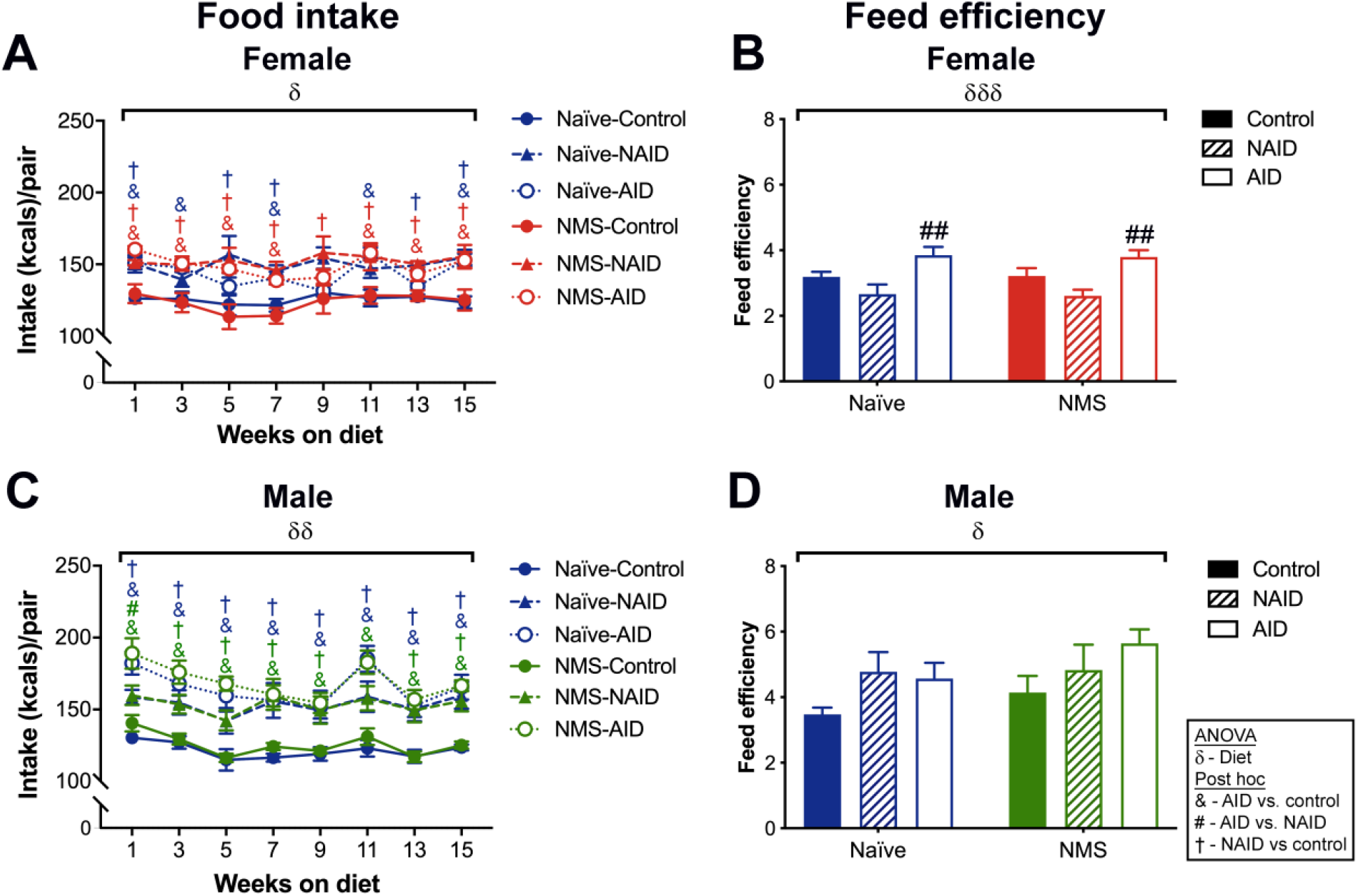
Effect of diet and early life stress on food intake and feed efficiency. **A**) In female mice, there was a significant effect of diet (*p*=0.042) on caloric intake. NMS and naïve mice fed either AID or NAID consumed more calories than their control-fed counterparts throughout the experiment. **B**) Feed efficiency in female mice was significantly impacted by diet (*p*=0.0004) with naïve-AID and NMS-AID mice having significantly higher feed efficiencies compared to their NAID-fed counterparts, despite being calorically identical. **C**) In male mice, there was a significant overall effect of diet (*p*=0.007) on caloric intake with AID- and NAID-fed mice consuming more calories than control-fed mice at every time point of the study. **D**) There was also a significant effect of diet (*p*=0.046) on feed efficiency in male mice, but no statistical difference between NAID- and AID-fed mice. δ denotes a significant effect of diet, three-way RM ANOVA (**A, C**) or two-way ANOVA (**B, C**). & *p*<0.05 AID vs. control, † *p*<0.05 NAID vs. control, #, ## *p*<0.05, 0.01 AID vs. NAID, Fisher’s LSD posttest. n=3-7 pairs per group.

### AID and NAID differentially affected serum glucose and insulin levels in male and female mice

Diet significantly impacted serum glucose levels during a glucose tolerance test (GTT), such that NMS-AID female mice had significantly higher serum glucose at 15, 30, and 60 minutes into the GTT (Figure 7A), and overall (Figure 7B) compared to NMS-control female mice. NMS-NAID female mice also had significantly higher GTT AUC measurements compared to NMS-control mice (Figure 7B). Female naïve-AID mice also had significantly higher serum glucose levels at 30 minutes into the GTT (Figure 7B) and overall (Figure 7C) compared to naïve-control female mice. No significant impact of NMS or diet was observed on fasting serum insulin levels (Figure 7C) or HOMA-IR measurements (Figure 7D) in female mice.

**Figure 7.**
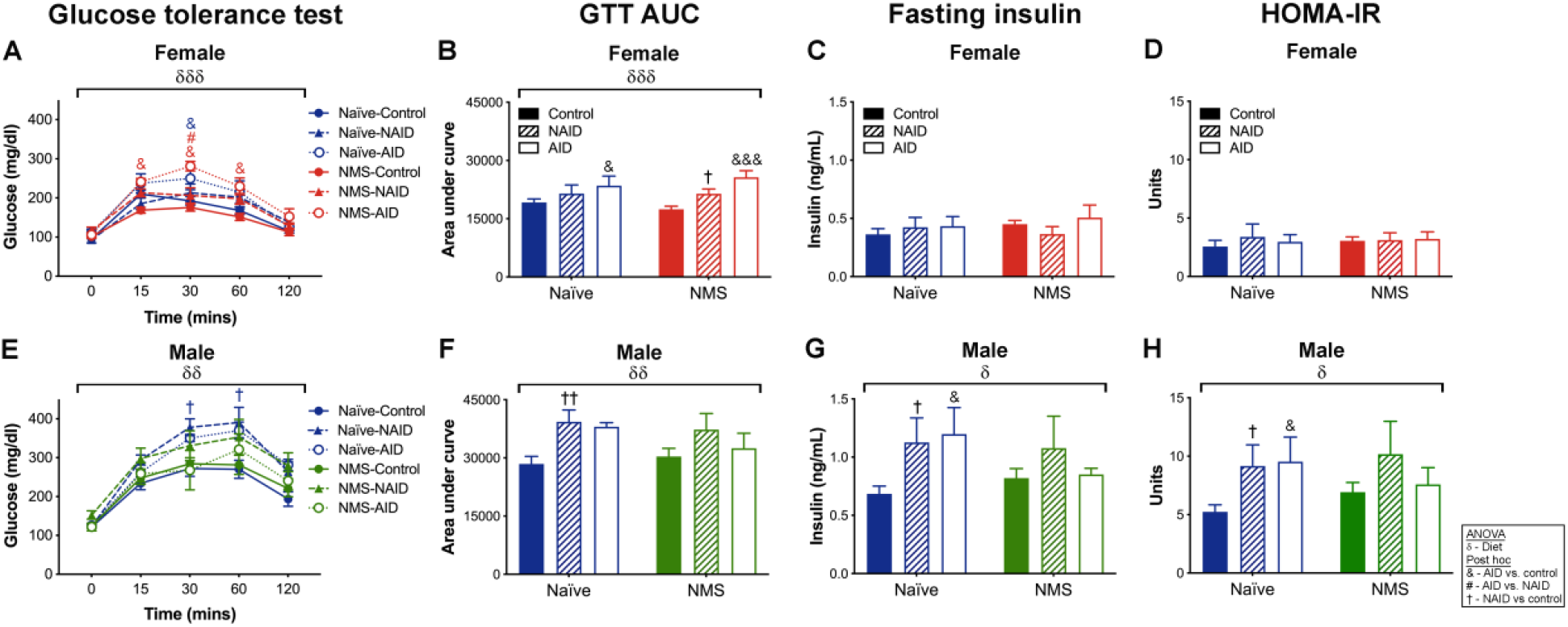
Impact of diet and early life stress on glucose tolerance, fasting insulin, and HOMA-IR. **A**) In female mice, there was a significant effect of diet (*p*<0.001) on glucose tolerance with female NMS-AID mice having significantly higher serum glucose compared to NMS-Control mice at 15, 30, and 60 minutes and higher than NMS-NAID mice at 30 minutes. Additionally, Naïve-AID mice had a higher serum glucose level at 30 minutes compared to Naïve-Control mice. **B**) Area under the curve (AUC) measurements revealed a significant diet effect (*p*=0.0003) with both naïve-AID and NMS-AID mice having significantly higher serum glucose compared to their control counterparts. NMS-NAID mice were also significantly higher than NMS-Control. There was no significant effect of diet or NMS on fasting serum insulin levels (**C**) or calculated HOMA-RI (**D**). **E**) In male mice, a significant effect of diet (*p*=0.004) was observed on glucose tolerance in male mice with naïve-NAID mice having significantly higher serum glucose levels at 30 and 60 minutes compared to naïve-control mice. **F**) AUC measurements were also significantly impacted by diet (*p*=0.005) with naïve-NAID mice having higher serum glucose levels than naïve-control mice. A significant impact of diet was observed on fasting serum insulin levels (*p*=0.040) and HOMA-IR (*p*=0.032) in male mice with naïve-AID and naïve-NAID mice having significantly higher levels than naïve-control mice. δ denotes a significant effect of diet, three-way RM ANOVA (**A, E**) or two-way ANOVA (**B-D, F-H**). &, &&& *p*<0.05, 0.001 AID vs. control, †, †† *p*<0.05, 0.01 NAID vs. control, # *p*<0.05 AID vs. NAID, Fisher’s LSD posttest. n=8-16 per group.

Diet significantly impacted serum glucose levels in male mice during a GTT, predominantly in naïve mice (Figure 7E-F). Naïve-NAID male mice had a significantly higher serum glucose level compared to male naïve-control mice at 30 and 60 minutes into the GTT (Figure 7E). There was a significant effect of diet on male GTT AUC with naïve-NAID male mice having significantly higher GTT AUC measurements compared to male naïve-control mice (Figure 7F). Diet also significantly impacted fasting serum insulin levels (Figure 7G) and HOMA-IR measurements in male mice (Figure 7H), such that naïve-NAID and naïve-AID mice were significantly higher than naïve-control mice. No significant diet effects were observed in NMS male mice. Overall, these data suggest that female NMS mice are more susceptible to NAID and AID compared to female naïve mice, whereas male mice, in general, have worsened outcomes due to NAID and AID, with naïve mice being more affected.

### Diet and NMS differentially affected serum corticosterone levels in female and male mice

Serum corticosterone levels were significantly impacted by diet in both female (Figure 8A) and male (Figure 8B) mice. Female NMS-AID mice had significantly lower serum corticosterone levels compared to female naïve-control mice (Figure 7-8A), whereas male NMS-AID mice had significantly higher serum corticosterone levels compared to NMS-control and NMS-NAID mice (Figure 8B). Male NMS-NAID mice also had significantly lower serum corticosterone levels compared to naïve-NAID mice. These results suggest that there is a profound sexual divergence in diet impact on corticosterone levels, particularly in response to AID.

**Figure 8.**
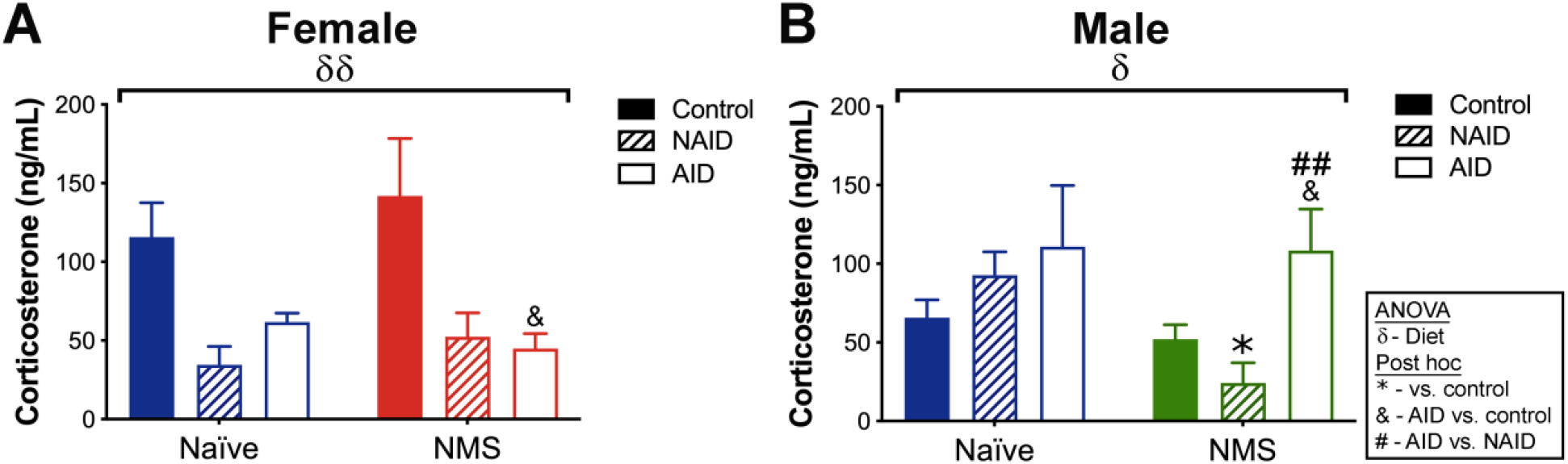
Effects of diet and early life stress on serum corticosterone concentrations. **A**) In female mice, there was a significant effect of diet (*p*=0.008) on serum corticosterone level. NMS-AID had significantly lower serum corticosterone compared to NMS-control mice. **B**) In male mice, there was also a significant effect of diet (*p*=0.021) on serum corticosterone levels. NMS-NAID mice had significantly lower serum corticosterone compared to naïve-NAID and NMS-AID mice and NMS-AID mice also had significantly higher corticosterone compared to NMS-control. δ denotes a significant effect of diet, two-way ANOVA. * *p*<0.05 vs. same diet-fed naïve, &, &&& *p*<0.05, 0.001 AID vs. control, ## *p*<0.01 AID vs. NAID, Fisher’s LSD posttest. n=8-16 per group.

### Diet and NMS significantly increased mRNA levels of macrophage and inflammatory markers in periovarian and epididymal adipose tissue

The mRNA levels of macrophage and inflammatory markers were measured in periovarian and epididymal fat pads (Figure 9) of female and male mice, respectively. In female mice, there was a significant effect of diet on the mRNA levels of F4/80 (general macrophage marker), CD68 (general macrophage marker), CD11b and CD11c (anti- and pro-inflammatory macrophage markers, respectively), and TNFα (pro-inflammatory cytokine) (Figure 9A). For nearly each gene, female AID-fed mice had significantly higher mRNA levels compared to control- and NAID-fed mice. In males, there was a significant effect of diet on CD68, CD11c, and IL-10 (anti-inflammatory cytokine) mRNA levels (Figure 9B). There was also a significant effect of NMS on CD11c mRNA levels. Male NMS-AID and -NAID-fed mice had significantly higher CD11c mRNA levels compared to NMS-control-fed mice and male naïve-AID-fed mice had a significantly higher IL-10 mRNA level compared to naïve-control-fed mice. These results suggest that AID largely drives periovarian adipose inflammatory marker expression in female mice, regardless of NMS status, whereas diet only impacts inflammatory markers in male NMS mice.

**Figure 9.**
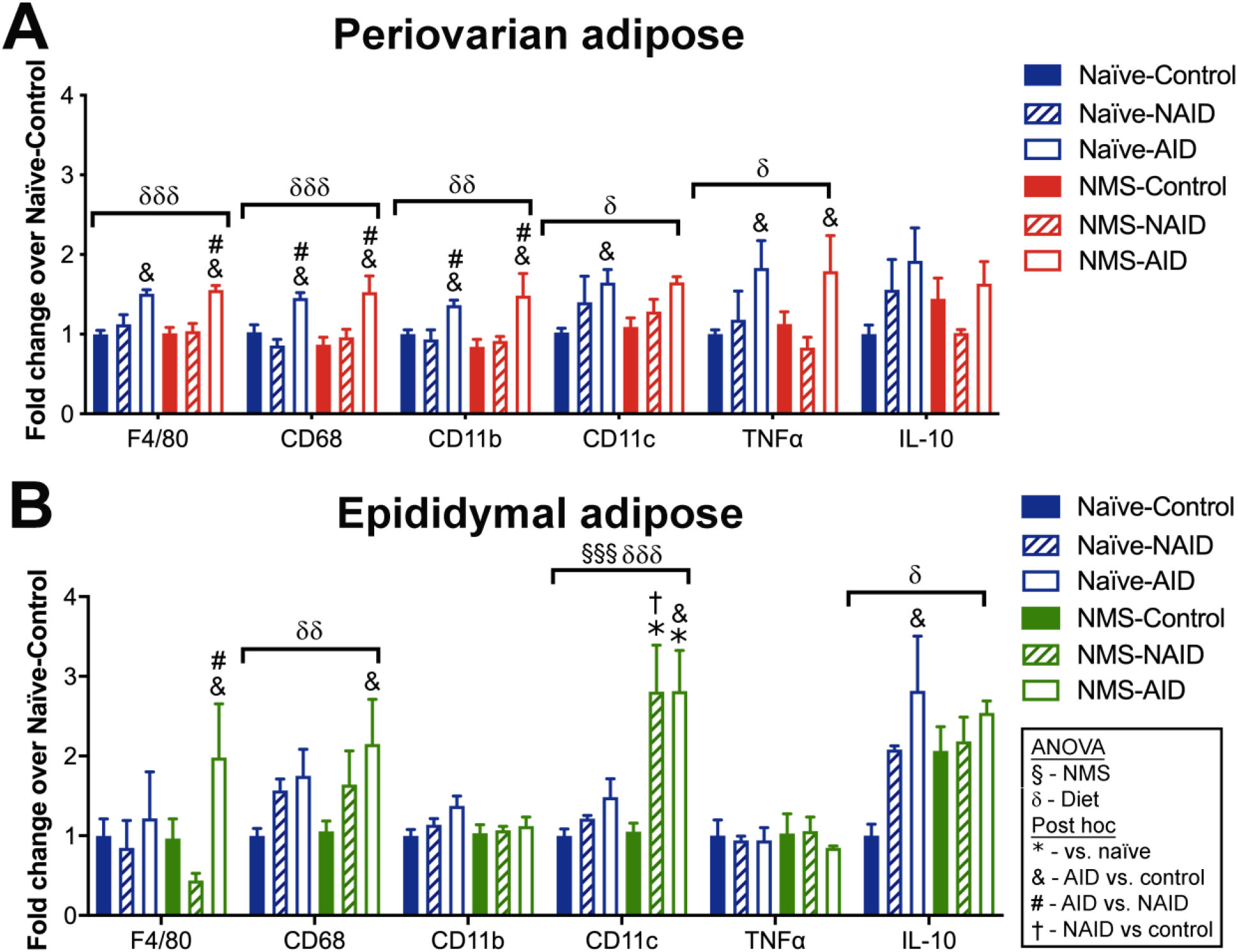
The effect of diet and early life stress on gene expression of inflammatory markers in gonadal adipose tissue. **A**) In females, there was a significant effect of diet on F4/80 (*p*<0.0001), CD68 (*p*<0.0001), CD11b (*p*=0.001), CD11c (*p*=0.024), and TNFα (*p*=0.016). AID-fed mice had significantly higher mRNA levels compared to both control-fed and NAID-fed for most genes analyzed, regardless of NMS status. **B**) In males, there was a significant effect of diet on CD68 (*p*=0.003), CD11c (*p*<0.0001), and IL-10 (*p*=0.021) and additional effects of NMS (*p*<0.0001) and an interaction effect of NMS and diet (*p*=0.012) on CD11c. F4/80 mRNA levels were significantly higher in NMS-AID mice compared to NMS-control or NMS-NAID mice. CD11c was significantly higher in NMS-AID and -NAID mice compared to NMS-control mice and to naïve mice fed the same diet. Il-10 was elevated in naïve-AID mice compared to naïve-Control mice. δ denotes a significant effect of diet, two-way ANOVA. * *p*<0.05 vs. same diet-fed naïve, & *p*<0.05 AID vs. control, † *p*<0.05 NAID vs. control, # *p*<0.05 AID vs. NAID, Fisher’s LSD posttest. n=5 per group.

## Discussion

Chronic pain is a highly prevalent and costly condition experienced by millions of Americans (1). It is associated with an increased inflammatory state and comorbid expression of other chronic pain disorders (39–43). In the present study, we used a model of NMS in mice to test whether an AID could attenuate early life stress-induced widespread hypersensitivity (30–32, 44, 45). We hypothesized that mice consuming an AID would be protected from developing NMS-induced urogenital and hindpaw hypersensitivity, compared to mice on a calorie and macronutrient matched NAID or a low fat control diet. Our results demonstrate a benefit of an AID on most behavioral outcomes, yet highlight the complex relationships between diet, stress, and sex on body weight and metabolic regulation, markers of inflammation, and corticosterone concentrations. Interestingly, the high-fat AID improved measures of hypersensitivity despite inducing increased adiposity and evidence of metabolic dysfunction, particularly in female mice.

Consumption of the AID increased perigenital withdrawal thresholds in all groups regardless of sex or early-life stress exposure. These findings are in line with studies which found improvements in pain outcomes following administration of individual anti-inflammatory compounds (16, 19, 21, 23, 24, 26, 27). Many of these studies (16, 19, 21, 26) were conducted in naïve, male mice with quantities of anti-inflammatory compounds that were 2-1000 times the amount used in the current study. Our study was unique in that it included both naïve and stressed conditions, female and male mice, and quantities of anti-inflammatory compounds that are realistically attainable in the human diet (46). There is evidence that this type of dietary intervention may translate well to humans as a recent systematic review (47) found that nutritional interventions significantly reduced pain scores, although the specific interventions were varied and additional high-quality clinical trials are needed. The inclusion of both sexes in this study led to some interesting albeit expected sex differences.

We found that only the female mice displayed NMS-induced hindpaw hypersensitivity suggesting that female mice are more susceptible to widespread sensitivity than male mice. It is widely accepted that this sex difference exists in the incidence of chronic pain disorders as well as in laboratory-evoked pain responses (48). Females are over represented in most pain conditions that present in both sexes including fibromyalgia, migraine, temporomandibular disorder, irritable bowel syndrome, and interstitial cystitis/painful bladder syndrome (49). Additionally, females exhibit increased sensitivity to evoked painful stimuli compared to males (50, 51). The mechanisms behind the sex differences in pain are attributed to a complex interaction between biological and psychosocial mechanisms. Preclinical work has revealed that pain processing is a complex circuitry that involves the peripheral and central nervous systems and both inhibitory and facilitatory mechanisms and sex hormones are known to influence this pathway at multiple levels through complicated interactions (52). This sex difference in hindpaw hypersensitivity is in line with previous findings and highlights the importance of studying both sexes in the field of pain research.

The high fat (35%) NAID elevated perigenital withdrawal thresholds in NMS male, but not female, mice which is similar to the increase in hindpaw allodynia that was previously found in males on 54% high fat diet (53). The anti-inflammatory components in the AID prevented this high fat diet-induced hypersensitivity, independent of body weight. These findings indicate that while females may generally be more susceptible to widespread sensitivity, the addition of high fat diet can reverse this sex difference and confer a greater susceptibility to allodynia in males, which coincides with the males’ increased risk of developing high fat diet-induced adverse metabolic effects.

Patients with chronic pain syndromes often experience symptoms of or are diagnosed with mood disorders, such as anxiety and depression (43, 54, 55). Anhedonia, or a lack of pleasure-seeking behavior, is a hallmark symptom of depression and can be measured behaviorally in rodents (56). Nest building is an innate and complex behavior that is impacted by hippocampal damage (36) and social defeat stress (57). The latter impact can be reversed by treatment with antidepressants, suggesting that nest building may be an appropriate measure for evaluating depressive-like behaviors. Here, we saw an overall diet effect on lowering nest building scores in the female, but not male, mice. Specifically, the female NMS mice on an AID showed a significantly lower nest score compared to naïve control-fed mice. We have previously reported on reduced regulatory gene expression in the hippocampus of NMS mice (29–32). These results suggest that AID may be exacerbating these deficits, thereby inducing a depressive-like state in these mice. Future studies are needed to fully explore this potential outcome.

It is well-known that males, compared to females, are more prone to the adverse health effects of high fat diet (58–60). Similar to our previous work using a 45% high fat diet (61), all male mice gained substantially more body weight and fat mass on the 35% high fat diets (AID and NAID) compared to the low fat control diet. This was largely driven by increased caloric intake although changes in energy expenditure and ambulatory cage activity were not measured and may also contribute to the phenotype. In line with this increased adiposity and body weight, markers of metabolic health including glucose tolerance and, in some cases, fasting insulin, were worsened by the AID and NAID in male mice.

An elevation in caloric intake was also seen in female mice fed either the AID or NAID, however, it was only the NMS-AID female mice that displayed elevated body weight and adiposity. This suggests that energy expenditure was affected by the added anti-inflammatory components, despite the negligible calorie content, and enhanced weight and adiposity gains following early life stress. This was surprising because the AID and NAID were derived from identical macronutrient sources/composition and energy density. It is also interesting that improvements in pain-like behaviors did not reduce weight gain or adiposity in NMS-AID-fed mice. This is similar to what was found in a different study using this AID, in which the AID-induced prevention of mechanical and thermal sensitivity in a model of inflammatory pain did not prevent increases body weight and adiposity (28). Additionally, an exercise intervention reversed high fat diet-induced mechanical allodynia and had no effect on body weight or glucose tolerance (53). There is a positive association between pain and body mass index (62) that is often assumed to be causally related. One prevailing hypothesis is that high levels of pain discourage physical activity, resulting in weight gain due to insufficient energy expenditure relative to energy intake. These data do not support this hypothesis but instead suggest that conditions that drive the development of chronic pain, such as NMS used in this study, may lead to long term adaptations in systems that regulate energy intake and expenditure in a manner that is independent of the development of long-term pain-like behaviors. It is worth noting that the circuitry regulating acute states of hunger and pain, overlap in complex ways that are only just beginning to be understood and may play a role in this model (63).

We have previously found altered corticosterone concentrations, a marker of HPA axis output, in this model of early life stress (29, 30). This is consistent with the clinical literature which shows both hypercortisolism (64, 65) and hypocortisolism (66, 67) in adults that report a history of childhood abuse or stress. We did not find a main effect of NMS in this cohort, although the measurement at the single timepoint (early in the light cycle) may have masked disruptions in the cyclicity of the circadian secretions or differences in peak concentrations. We did find a sex-specific main effect of diet in which the female mice on either of the diets high in fat (AID and NAID) displayed reduced corticosterone concentrations compared to control diet-fed females. The reason for this decrease is unclear, although it is known that high fat diet can decrease adipose tissue 11 β-hydroxysteroid dehydrogenase type 1 (11 β-HSD1) (68, 69), which converts inactive corticosterone into active corticosterone (70). Additionally, its known that high fat diet, through endogenous opioid secretion (71, 72), can dampen HPA axis activation (73). Both of these mechanisms would theoretically decrease the amount of circulating corticosterone; however, future studies are needed to determine the contribution of either of these mechanisms to this phenotype. It is unclear why the females consistently saw this diet effect while the males did not, although sex differences in the HPA axis including corticosterone secretion and availability are widely known and influenced by the sex hormones (74–77).

Many chronic pain disorders are associated with a chronic state of low-grade inflammation (4, 78, 79) and an AID or diet supplemented with anti-inflammatory substances has been shown to attenuate or prevent the development and pain in humans (80) and rodents (81–84). We expected the AID-induced improvements in pain-like behaviors to coincide with reductions in tissue inflammation. Not only did the gene expression profile of pro-inflammatory markers within the gonadal fat pad not show reductions but, in some cases, levels were elevated in the NMS mice (F4/80 (females) and CD68 (males and females)). These findings may have been complicated by the increased adiposity and visceral fat of the NMS-AID-fed mice as visceral adipose accumulation is associated with chronic low-grade inflammation (9, 85, 86). Although we did not measure inflammation in other peripheral or central tissues, it is possible there were improvements in inflammation in those tissues that contributed to improvements in the widespread hypersensitivity independent of the inflammation in adipose tissue depots. Future studies could benefit from preventing the increased adiposity by restricting intake on the AID or the addition of a low fat diet supplemented with the anti-inflammatory compounds to determine how an AID affects inflammation in the absence of increased adiposity.

### Conclusions

In summary, the AID improved perigenital withdrawal thresholds in all mice regardless of sex or early life stress exposure, and selectively improved hindpaw allodynia in female NMS mice. This improvement in pain-like behaviors was seen despite AID-induced increases in body weight and adiposity, glucose intolerance, and pro-inflammatory markers. Both the AID and NAID increased body weight and adiposity suggesting that, at least in this preclinical model, the high fat diet-induced increase in food intake was metabolically detrimental and was not overcome by the inclusion of anti-inflammatory compounds which, in the case of the NMS females, worsened weight gain and adiposity. If these findings translate to clinical populations, it may suggest that there are limits to the overall benefit of an anti-inflammatory diet if total intake is not limited.

## References

1. Dahlhamer J, Lucas J, Zelaya C, Nahin R, Mackey S, DeBar L, Kerns R, Von Korff M, Porter L, Helmick C. (2018) Prevalence of chronic pain and high-impact chronic pain among adults - united states, 2016. MMWR Morb Mortal Wkly Rep. 67:(36):1001–1006.

2. Pitcher MH, Von Korff M, Bushnell MC, Porter L. (2019) Prevalence and profile of high-impact chronic pain in the united states. J Pain. 20(2): 146–160.

3. Gaskin DJ, Richard P. (2012) The economic costs of pain in the united states. J Pain. 13(8):715–24.

4. Zhang JM, An J. (2007) Cytokines, inflammation, and pain. Int Anesthesiol Clin. 45(2):27–37.

5. Abbadie C, Bhangoo S, De Koninck Y, Malcangio M, Melik-Parsadaniantz S, White FA. (2009) Chemokines and pain mechanisms. Brain Res Rev. 60(1): 125–34.

6. Ren K, Dubner R. (2010) Interactions between the immune and nervous systems in pain. Nat Med. 16(11):1267–76.

7. Schaible HG, von Banchet GS, Boettger MK, Brauer R, Gajda M, Richter F, Hensellek S, Brenn D, Natura G. (2010) The role of proinflammatory cytokines in the generation and maintenance of joint pain. Ann N Y Acad Sci. 1193:60–9.

8. Gold MS, Gebhart GF. (2010) Nociceptor sensitization in pain pathogenesis. Nat Med. 16(11):1248–57.

9. Gregor MF, Hotamisligil GS. (2011) Inflammatory mechanisms in obesity. Annu Rev Immunol. 29:415–45.

10. Eichwald T, Talbot S. (2020) Neuro-immunity controls obesity-induced pain. Front Hum Neurosci. 14:181.

11. Kidd BL, Urban LA. (2001) Mechanisms of inflammatory pain. Br J Anaesth. 87(1):3–11.

12. Payne R. (2000) Limitations of nsaids for pain management: Toxicity or lack of efficacy? J Pain. 1(3 Suppl):14–8.

13. Tick H. (2015) Nutrition and pain. Phys Med Rehabil Clin N Am. 26(2):309–20.

14. Ricker MA, Haas WC. (2017) Anti-inflammatory diet in clinical practice: A review. Nutr Clin Pract. 32(3):318–325.

15. Higdon JV, Frei B. (2003) Tea catechins and polyphenols: Health effects, metabolism, and antioxidant functions. Crit Rev Food Sci Nutr. 43(1):89–143.

16. Kuang X, Huang Y, Gu HF, Zu XY, Zou WY, Song ZB, Guo QL. (2012) Effects of intrathecal epigallocatechin gallate, an inhibitor of toll-like receptor 4, on chronic neuropathic pain in rats. Eur J Pharmacol. 676(1-3):51–6.

17. Guerrero-Beltran CE, Calderon-Oliver M, Pedraza-Chaverri J, Chirino YI. (2012) Protective effect of sulforaphane against oxidative stress: Recent advances. Exp Toxicol Pathol. 64(5):503–8.

18. Lee S, Kim J, Seo SG, Choi BR, Han JS, Lee KW, Kim J. (2014) Sulforaphane alleviates scopolamine-induced memory impairment in mice. Pharmacol Res. 85:23–32.

19. Wang C, Wang C. (2017) Anti-nociceptive and anti-inflammatory actions of sulforaphane in chronic constriction injury-induced neuropathic pain mice. Inflammopharmacology. 25(1):99–106.

20. Rocha-Gonzalez HI, Ambriz-Tututi M, Granados-Soto V. (2008) Resveratrol: A natural compound with pharmacological potential in neurodegenerative diseases. CNS Neurosci Ther. 14(3):234–47.

21. Sharma S, Kulkarni SK, Chopra K. (2007) Effect of resveratrol, a polyphenolic phytoalexin, on thermal hyperalgesia in a mouse model of diabetic neuropathic pain. Fundam Clin Pharmacol. 21(1):89–94.

22. Strimpakos AS, Sharma RA. (2008) Curcumin: Preventive and therapeutic properties in laboratory studies and clinical trials. Antioxid Redox Signal. 10(3):511–45.

23. Kuptniratsaikul V, Dajpratham P, Taechaarpornkul W, Buntragulpoontawee M, Lukkanapichonchut P, Chootip C, Saengsuwan J, Tantayakom K, Laongpech S. (2014) Efficacy and safety of curcuma domestica extracts compared with ibuprofen in patients with knee osteoarthritis: A multicenter study. Clin Interv Aging. 9:451–8.

24. Di YX, Hong C, Jun L, Renshan G, Qinquan L. (2014) Curcumin attenuates mechanical and thermal hyperalgesia in chronic constrictive injury model of neuropathic pain. Pain Ther. 3(1):59–69.

25. Zhang L, Virgous C, Si H. (2017) Ginseng and obesity: Observations and understanding in cultured cells, animals and humans. J Nutr Biochem. 44:1–10.

26. Nah JJ, Hahn JH, Chung S, Choi S, Kim YI, Nah SY. (2000) Effect of ginsenosides, active components of ginseng, on capsaicin-induced pain-related behavior. Neuropharmacology. 39(11):2180–4.

27. Silva AR, Bernardo A, Costa J, Cardoso A, Santos P, de Mesquita MF, Vaz Patto J, Moreira P, Silva ML, Padrao P. (2019) Dietary interventions in fibromyalgia: A systematic review. Ann Med. 51(sup1):2–14.

28. Totsch SK, Meir RY, Quinn TL, Lopez SA, Gower BA, Sorge RE. (2018) Effects of a standard american diet and an anti-inflammatory diet in male and female mice. Eur J Pain. 22(7):1203–1213.

29. Pierce AN, Eller-Smith OC, Christianson JA. (2018) Voluntary wheel running attenuates urinary bladder hypersensitivity and dysfunction following neonatal maternal separation in female mice. Neurourol Urodyn. 37(5):1623–1632.

30. Fuentes IM, Pierce AN, Di Silvestro ER, Maloney MO, Christianson JA. (2017) Differential influence of early life and adult stress on urogenital sensitivity and function in male mice. Front Syst Neurosci. 11:97.

31. Pierce AN, Di Silvestro ER, Eller OC, Wang R, Ryals JM, Christianson JA. (2016) Urinary bladder hypersensitivity and dysfunction in female mice following early life and adult stress. Brain Res. 1639:58–73.

32. Pierce AN, Ryals JM, Wang R, Christianson JA. (2014) Vaginal hypersensitivity and hypothalamic-pituitary-adrenal axis dysfunction as a result of neonatal maternal separation in female mice. Neuroscience. 263:216–30.

33. Eller OC, Morris EM, Thyfault JP, Christianson JA. (2020) Early life stress reduces voluntary exercise and its prevention of diet-induced obesity and metabolic dysfunction in mice. Physiol Behav. 223:113000.

34. Chaplan SR, Bach FW, Pogrel JW, Chung JM, Yaksh TL. (1994) Quantitative assessment of tactile allodynia in the rat paw. J Neurosci Methods. 53(1):55–63.

35. Dixon WJ. (1980) Efficient analysis of experimental observations. Annu Rev Pharmacol Toxicol. 20:441–62.

36. Deacon RM, Croucher A, Rawlins JN. (2002) Hippocampal cytotoxic lesion effects on species-typical behaviours in mice. Behav Brain Res. 132(2):203–13.

37. Deacon R. (2012) Assessing burrowing, nest construction, and hoarding in mice. J Vis Exp (59):e2607.

38. Pfaffl MW. (2001) A new mathematical model for relative quantification in real-time rt-pcr. Nucleic Acids Res. 29(9):e45.

39. Clemens JQ, Meenan RT, O’Keeffe Rosetti MC, Kimes TA, Calhoun EA. (2008) Casecontrol study of medical comorbidities in women with interstitial cystitis. J Urol. 179(6):2222–5.

40. Arnold LD, Bachmann GA, Rosen R, Kelly S, Rhoads GG. (2006) Vulvodynia: Characteristics and associations with comorbidities and quality of life. Obstet Gynecol. 107(3):617–24.

41. Rodriguez MA, Afari N, Buchwald DS, National Institute of D, Digestive, Kidney Diseases Working Group on Urological Chronic Pelvic P. (2009) Evidence for overlap between urological and nonurological unexplained clinical conditions. J Urol. 182(5):2123–31.

42. Aaron LA, Buchwald D. (2001) A review of the evidence for overlap among unexplained clinical conditions. Ann Intern Med. 134(9 Pt 2):868–81.

43. Clemens JQ, Brown SO, Calhoun EA. (2008) Mental health diagnoses in patients with interstitial cystitis/painful bladder syndrome and chronic prostatitis/chronic pelvic pain syndrome: A case/control study. J Urol. 180(4):1378–82.

44. Fuentes IM, Pierce AN, O’Neil PT, Christianson JA. (2015) Assessment of perigenital sensitivity and prostatic mast cell activation in a mouse model of neonatal maternal separation. J Vis Exp (102):e53181.

45. Fuentes IM, Christianson JA. (2018) The influence of early life experience on visceral pain. Front Syst Neurosci. 12:2.

46. Totsch SK, Waite ME, Sorge RE. (2015) Dietary influence on pain via the immune system. Prog Mol Biol Transl Sci. 131:435–69.

47. Brain K, Burrows TL, Rollo ME, Chai LK, Clarke ED, Hayes C, Hodson FJ, Collins CE. (2019) A systematic review and meta-analysis of nutrition interventions for chronic noncancer pain. J Hum Nutr Diet. 32(2):198–225.

48. Sorge RE, Totsch SK. (2017) Sex differences in pain. J Neurosci Res. 95(6):1271–1281.

49. Berkley KJ. (1997) Sex differences in pain. Behav Brain Sci. 20(3):371-80; discussion 435-513.

50. Fillingim RB. (2000) Sex, gender, and pain: Women and men really are different. Curr Rev Pain. 4(1):24–30.

51. Mogil JS. (2012) Sex differences in pain and pain inhibition: Multiple explanations of a controversial phenomenon. Nat Rev Neurosci. 13(12):859–66.

52. Fillingim RB, Ness TJ. (2000) Sex-related hormonal influences on pain and analgesic responses. Neurosci Biobehav Rev. 24(4):485–501.

53. Cooper MA, Ryals JM, Wu PY, Wright KD, Walter KR, Wright DE. (2017) Modulation of diet-induced mechanical allodynia by metabolic parameters and inflammation. J Peripher Nerv Syst. 22(1):39–46.

54. Demyttenaere K, Bruffaerts R, Lee S, Posada-Villa J, Kovess V, Angermeyer MC, Levinson D, de Girolamo G, Nakane H, Mneimneh Z, Lara C, de Graaf R, Scott KM, Gureje O, Stein DJ, Haro JM, Bromet EJ, Kessler RC, Alonso J, Von Korff M. (2007) Mental disorders among persons with chronic back or neck pain: Results from the world mental health surveys. Pain. 129(3):332–42.

55. Gureje O, Von Korff M, Kola L, Demyttenaere K, He Y, Posada-Villa J, Lepine JP, Angermeyer MC, Levinson D, de Girolamo G, Iwata N, Karam A, Guimaraes Borges GL, de Graaf R, Browne MO, Stein DJ, Haro JM, Bromet EJ, Kessler RC, Alonso J. (2008) The relation between multiple pains and mental disorders: Results from the world mental health surveys. Pain. 135(1-2):82–91.

56. Schmidt MV, Wang XD, Meijer OC. (2011) Early life stress paradigms in rodents: Potential animal models of depression? Psychopharmacology (Berl). 214(1):131–40.

57. Otabi H, Goto T, Okayama T, Kohari D, Toyoda A. (2017) The acute social defeat stress and nest-building test paradigm: A potential new method to screen drugs for depressive-like symptoms. Behav Processes. 135:71–75.

58. Gelineau RR, Arruda NL, Hicks JA, Monteiro De Pina I, Hatzidis A, Seggio JA. (2017) The behavioral and physiological effects of high-fat diet and alcohol consumption: Sex differences in c57bl6/j mice. Brain Behav. 7(6):e00708.

59. Hwang LL, Wang CH, Li TL, Chang SD, Lin LC, Chen CP, Chen CT, Liang KC, Ho IK, Yang WS, Chiou LC. (2010) Sex differences in high-fat diet-induced obesity, metabolic alterations and learning, and synaptic plasticity deficits in mice. Obesity (Silver Spring). 18(3):463–9.

60. Foright RM, Johnson GC, Kahn D, Charleston CA, Presby DM, Bouchet CA, Wellberg EA, Sherk VD, Jackman MR, Greenwood BN, MacLean PS. (2020) Compensatory eating behaviors in male and female rats in response to exercise training. Am J Physiol Regul Integr Comp Physiol.

61. Eller OC, Morris EM, Thyfault JP, Christianson JA. (2020) Early life stress reduces voluntary exercise and its prevention of diet-induced obesity and metabolic dysfunction in mice. Physiol Behav:113000.

62. Stone AA, Broderick JE. (2012) Obesity and pain are associated in the united states. Obesity (Silver Spring). 20(7):1491–5.

63. Alhadeff AL, Su Z, Hernandez E, Klima ML, Phillips SZ, Holland RA, Guo C, Hantman AW, De Jonghe BC, Betley JN. (2018) A neural circuit for the suppression of pain by a competing need state. Cell. 173(1):140–152 e15.

64. Heim C, Newport DJ, Bonsall R, Miller AH, Nemeroff CB. (2001) Altered pituitary-adrenal axis responses to provocative challenge tests in adult survivors of childhood abuse. Am J Psychiatry. 158(4):575–81.

65. Tyrka AR, Wier L, Price LH, Ross N, Anderson GM, Wilkinson CW, Carpenter LL. (2008) Childhood parental loss and adult hypothalamic-pituitary-adrenal function. Biol Psychiatry. 63(12):1147–54.

66. Heim C, Ehlert U, Hanker JP, Hellhammer DH. (1998) Abuse-related posttraumatic stress disorder and alterations of the hypothalamic-pituitary-adrenal axis in women with chronic pelvic pain. Psychosom Med. 60(3):309–18.

67. Gunnar MR, Quevedo KM. (2008) Early care experiences and hpa axis regulation in children: A mechanism for later trauma vulnerability. Prog Brain Res. 167:137–49.

68. Morton NM, Ramage L, Seckl JR. (2004) Down-regulation of adipose 11beta-hydroxysteroid dehydrogenase type 1 by high-fat feeding in mice: A potential adaptive mechanism counteracting metabolic disease. Endocrinology. 145(6):2707–12.

69. Drake AJ, Livingstone DE, Andrew R, Seckl JR, Morton NM, Walker BR. (2005) Reduced adipose glucocorticoid reactivation and increased hepatic glucocorticoid clearance as an early adaptation to high-fat feeding in wistar rats. Endocrinology. 146(2):913–9.

70. Tomlinson JW, Walker EA, Bujalska IJ, Draper N, Lavery GG, Cooper MS, Hewison M, Stewart PM. (2004) 11beta-hydroxysteroid dehydrogenase type 1: A tissue-specific regulator of glucocorticoid response. Endocr Rev. 25(5):831–66.

71. Tsujii S, Nakai Y, Fukata J, Nakaishi S, Takahashi H, Usui T, Imura H. (1987) Effects of food deprivation and high fat diet on immunoreactive beta-endorphin levels in brain regions of zucker rats. Endocrinol Jpn. 34(6):903–9.

72. Tsujii S, Nakai Y, Fukata J, Nakaishi S, Takahashi H, Usui T, Imura H. (1987) Effects of food deprivation and high fat diet on immunoreactive dynorphin a(1-8) levels in brain regions of zucker rats. Peptides. 8(6):1075–8.

73. Drolet G, Dumont EC, Gosselin I, Kinkead R, Laforest S, Trottier JF. (2001) Role of endogenous opioid system in the regulation of the stress response. Prog Neuropsychopharmacol Biol Psychiatry. 25(4):729–41.

74. Kalinichev M, Easterling KW, Plotsky PM, Holtzman SG. (2002) Long-lasting changes in stress-induced corticosterone response and anxiety-like behaviors as a consequence of neonatal maternal separation in long-evans rats. Pharmacol Biochem Behav. 73(1):131–40.

75. McCormick CM, Robarts D, Kopeikina K, Kelsey JE. (2005) Long-lasting, sex-and age-specific effects of social stressors on corticosterone responses to restraint and on locomotor responses to psychostimulants in rats. Horm Behav. 48(1):64–74.

76. Wigger A, Neumann ID. (1999) Periodic maternal deprivation induces gender-dependent alterations in behavioral and neuroendocrine responses to emotional stress in adult rats. Physiol Behav. 66(2):293–302.

77. Kudielka BM, Kirschbaum C. (2005) Sex differences in hpa axis responses to stress: A review. Biol Psychol. 69(1):113–32.

78. Scarpellini E, Tack J. (2012) Obesity and metabolic syndrome: An inflammatory condition. Dig Dis. 30(2):148–53.

79. Miller AH, Raison CL. (2016) The role of inflammation in depression: From evolutionary imperative to modern treatment target. Nat Rev Immunol. 16(1):22–34.

80. Keske MA, Ng HL, Premilovac D, Rattigan S, Kim JA, Munir K, Yang P, Quon MJ. (2015) Vascular and metabolic actions of the green tea polyphenol epigallocatechin gallate. Curr Med Chem. 22(1):59–69.

81. Bose M, Lambert JD, Ju J, Reuhl KR, Shapses SA, Yang CS. (2008) The major green tea polyphenol, (-)-epigallocatechin-3-gallate, inhibits obesity, metabolic syndrome, and fatty liver disease in high-fat-fed mice. J Nutr. 138(9):1677–83.

82. Choi KM, Lee YS, Kim W, Kim SJ, Shin KO, Yu JY, Lee MK, Lee YM, Hong JT, Yun YP, Yoo HS. (2014) Sulforaphane attenuates obesity by inhibiting adipogenesis and activating the ampk pathway in obese mice. J Nutr Biochem. 25(2):201–7.

83. Jeon BT, Jeong EA, Shin HJ, Lee Y, Lee DH, Kim HJ, Kang SS, Cho GJ, Choi WS, Roh GS. (2012) Resveratrol attenuates obesity-associated peripheral and central inflammation and improves memory deficit in mice fed a high-fat diet. Diabetes. 61(6):1444–54.

84. Weisberg SP, Leibel R, Tortoriello DV. (2008) Dietary curcumin significantly improves obesity-associated inflammation and diabetes in mouse models of diabesity. Endocrinology. 149(7):3549–58.

85. Lumeng CN, Deyoung SM, Bodzin JL, Saltiel AR. (2007) Increased inflammatory properties of adipose tissue macrophages recruited during diet-induced obesity. Diabetes. 56(1):16–23.

86. Weisberg SP, McCann D, Desai M, Rosenbaum M, Leibel RL, Ferrante AW, Jr. (2003) Obesity is associated with macrophage accumulation in adipose tissue. J Clin Invest. 112(12):1796–808.

